# Segment AnyNeuron

**DOI:** 10.1101/2024.08.24.609505

**Authors:** Taha Razzaq, Ahmed Qazi, Asim Iqbal

**Affiliations:** Tibbling Technologies, Redmond, WA, USA

## Abstract

Image segmentation plays an integral part in neuroimage analysis and is crucial for understanding brain disorders. Deep Learning (DL) models have shown exponential success in computer vision tasks over the years, including image segmentation. However, to achieve optimal performance, DL models require extensive annotated data for training, which is often the bottleneck to expediting brain-wide image analysis. For segmenting cellular structures such as neurons, the annotation process is cumbersome and time-consuming due to the inherent structural, intensity, and background variations present in the data caused by genetic markers, imaging techniques, etc. We propose an Active Learning-based neuron segmentation framework (Segment AnyNeuron), which incorporates state-of-the-art image segmentation modules - Detectron2 and HQ SAM, and requires minimal ground truth annotation to achieve high precision for brain-wide segmentation of neurons. Our framework can classify and segment completely unseen neuronal data by selecting the most representative samples for manual annotation, thus avoiding the cold-start problem common in Active Learning. We demonstrate the effectiveness of our framework for automated brain-wide segmentation of neurons on a variety of open-source neuron imaging datasets, acquired from different scanners and a variety of transgenic mouse lines.

## Introduction

Recent advancements in Deep Learning (DL) have revolutionized computer vision, demonstrating tremendous success in tasks such as object detection (Wu et al. [2019]) and image segmentation (Ke et al. [2023]). However, despite these successes, a significant challenge persists: DL models require large quantities of annotated data for training, which often proves to be a bottleneck (Minaee et al. [2021]). When sufficient labeled data is available, DL models for image segmentation and object detection exhibit remarkable performance on downstream tasks and are actively utilized in medical image analysis (Ronneberger et al. [2015a]). While DL offers substantial benefits for numerous medical applications, including disease diagnosis, treatment planning, and biological research, the requirement for extensive data remains a limiting factor due to the high cost and time involved in annotation (Ronneberger et al. [2015a]). This challenge is particularly pronounced in neuron segmentation, where the small and intricate structures make manual annotation exceptionally laborious and time-consuming.

To address this issue, Active Learning (AL) is a widely adopted approach designed to minimize the time and resources required for manual annotation (Mittal et al. [2023]). AL strategically selects the most representative and informative samples from a pool of unlabelled data. These samples, once manually annotated, are used to train or finetune the model, yielding significantly better results in less time. Given that AL is a well-studied solution in the context of image segmentation (Mackowiak et al. [2018]) and object detection (Kao et al. [2018]), it is increasingly being leveraged to enhance deep learning models in medical imaging (Nath et al. [2021]). Although AL-based medical image segmentation models have seen significant advancements over the past few years (Boehringer et al. [2023] et al. [2021]), the application of AL to neuron segmentation remains rare (Iqbal et al. [2019]).

We propose a novel Active Learning-based framework for neuron segmentation that leverages state-ofthe-art (SOTA) image detection and segmentation models, specifically Detectron2 (Wu et al. [2019]) and HQ-SAM (Ke et al. [2023]). While Detectron2 is one of the most commonly used detection models for medical images (Salh and Ali [2024])(Ali et al. [2022])(Chincholi and Koestler [2023]), HQ-SAM is also being integrated into medical applications (Zhang et al. [2024]). Our framework is designed to operate with minimal ground truth annotation, significantly reducing the annotation burden while maintaining high performance on unseen neuronal data. The key innovation of our approach lies in the integration of instance detection and Active Learning to iteratively refine the centers of the outputted bounding boxes (key points) and enhance segmentation accuracy. By using Detectron2 to generate initial key points on unseen, unlabelled data, we provide a strong baseline that can be corrected with minimal manual intervention. These corrected key points are then used by HQ-SAM to generate precise segmentation masks. Additionally, our framework includes an intensity-based thresholding feature that allows users to control the segmentation output by adjusting the intensity of detected neurons, providing flexibility and customization based on specific requirements. Our methodology incorporates advanced preprocessing steps such as intensity normalization and patch-based image segmentation, ensuring that our model receives the cleanest and most relevant data inputs. We demonstrate the effectiveness of our approach through analyses of a disease dataset, showcasing its adaptability and superior performance compared to existing methods. We aim to open-source our framework and provide a comprehensive guide on applying our Active Learning framework to novel datasets.

## Results & Discussion

We propose Segment AnyNeuron, a multi-step framework designed to optimize segmentation performance on novel neuron data. The framework consists of a neuron detector and segmenter, which, in conjunction with the Active Learning module, deliver benchmarking performance on unlabeled neuron datasets with minimal manual annotation. The overall pipeline of our framework, Segment AnyNeuron, is shown in **Figure 1**.

**Figure 1:**
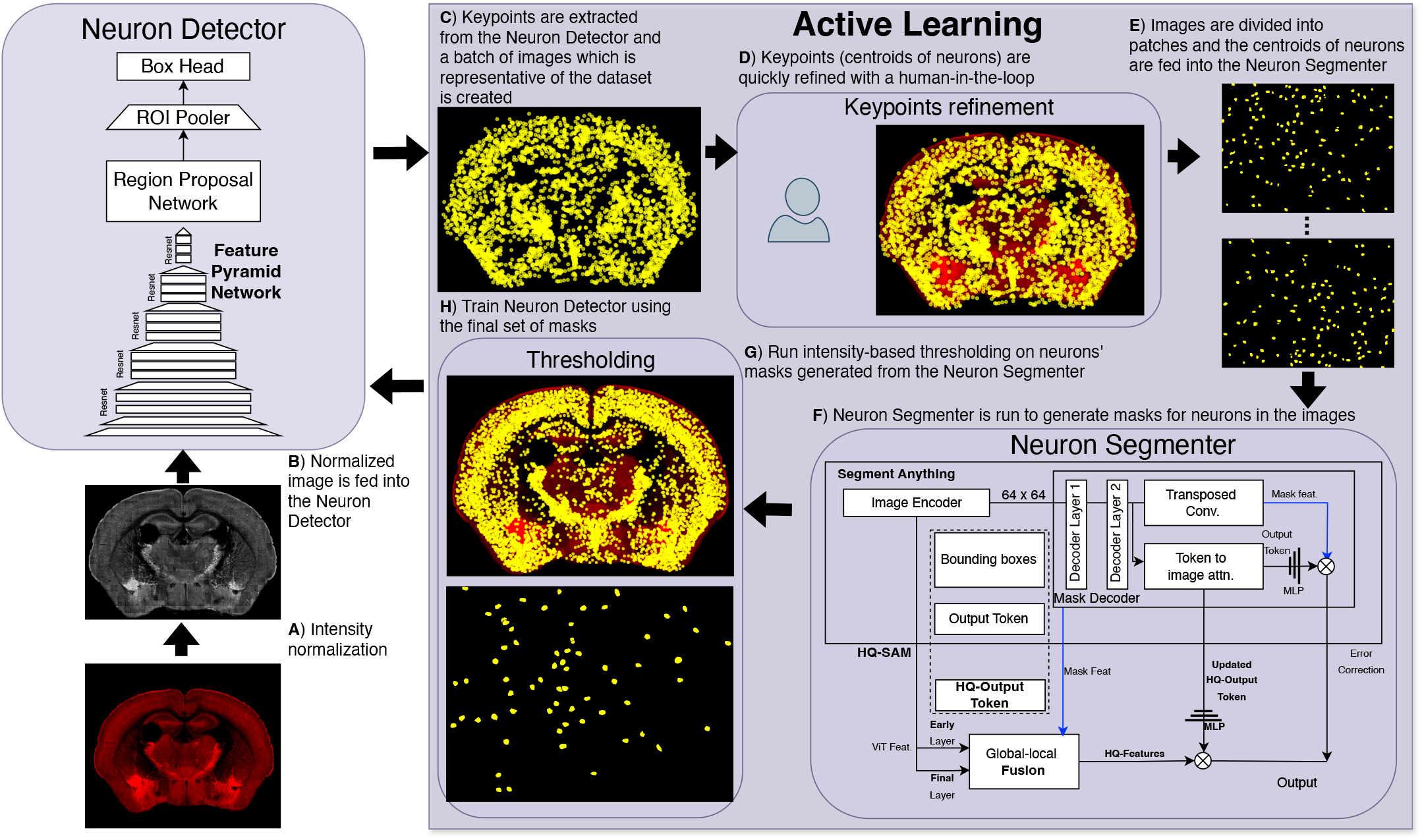
Block diagram for Segment AnyNeuron. A,B) An intensity-normalized version of the input unlabelled image is generated and fed into the Neuron Detector to generate keypoints (dirty ground truth). C) Representative samples from the entire unlabelled dataset are selected and fed into the Active Learning pipeline. D,E) After manual annotation fixing, the refined keypoints are F) processed by the neuron segmenter to generate masks, which are further G) refined through thresholding. H) The data is then used to train/finetune the Neuron Detector.

### Image datasets

#### SOM-Cre mouse line dataset

To evaluate the performance of our Active Learning pipeline, we employ open-source data from the Allen Brain Data Repository, focusing on transgenic somatostatin-Cre (SOM-Cre) mouse strains (Sst-IRES-Cre;Ai14). The SOM-Cre strain is extensively studied to elucidate the physiological role of somatostatin-expressing neurons in the mouse brain (Viollet et al. [2017]) and its association with Alzheimer’s disease (Williams et al. [2023 o et al. [2021]), thus providing a pertinent dataset for our experimental validation. The dataset comprises detailed neuronal mouse brain sections, from which we strategically select a few for our Active Learning pipeline.

#### Fluocells v2 dataset

Since our (pre-trained) model was primarily fine-tuned on an in-house neuron dataset, we sought to demonstrate its effectiveness on similar datasets by selecting the Fluorescent Neuronal Cells v2 dataset. Fluocells comprise three fluorescence microscopy image collections, where rodent neuronal cell nuclei and cytoplasm are stained with cFOS, the b-subunit of Cholera Toxin (CTb), and orexin markers, highlighting their anatomical and functional characteristics. Ground truth annotations for these images are publicly available. Prior to model input, all images underwent standard pre-processing procedures, including intensity normalization and patching.

#### Active Learning performance on SOM-Cre mouse line dataset

To evaluate the effectiveness of our proposed Active Learning pipeline, we utilize unseen SOM-Cre mouse line samples from the Allen Brain Data repository. Given the differences in neuron size and structure compared to our initial training data, the current model checkpoint demonstrates suboptimal performance on this novel dataset. To address this, we implement Active Learning. Our neuron detector generates preliminary (dirty) ground truth for a strategically chosen subset of samples, which are then manually corrected. These corrected samples, along with their key points, are used by the segmentation model to generate the corresponding masks. The generated masks are further refined using an intensity thresholding parameter, which enables the elimination of extraneous neurons, thereby producing a more accurate and cleaner ground truth mask. After pre-processing, the images and their masks are converted into patches and fed into our model for fine-tuning. We conduct minimal fine-tuning (approximately 10 epochs) and present the qualitative results of our model before and after Active Learning as seen in **Figure 2**. It presents distinct sections of the mouse brain, accompanied by the masks generated by our pipeline before and after the application of Active Learning. In addition to the full section masks, the top rows (1*st* and 3*rd* rows) display zoomed-in subsections with their corresponding masks overlaid.

**Figure 2:**
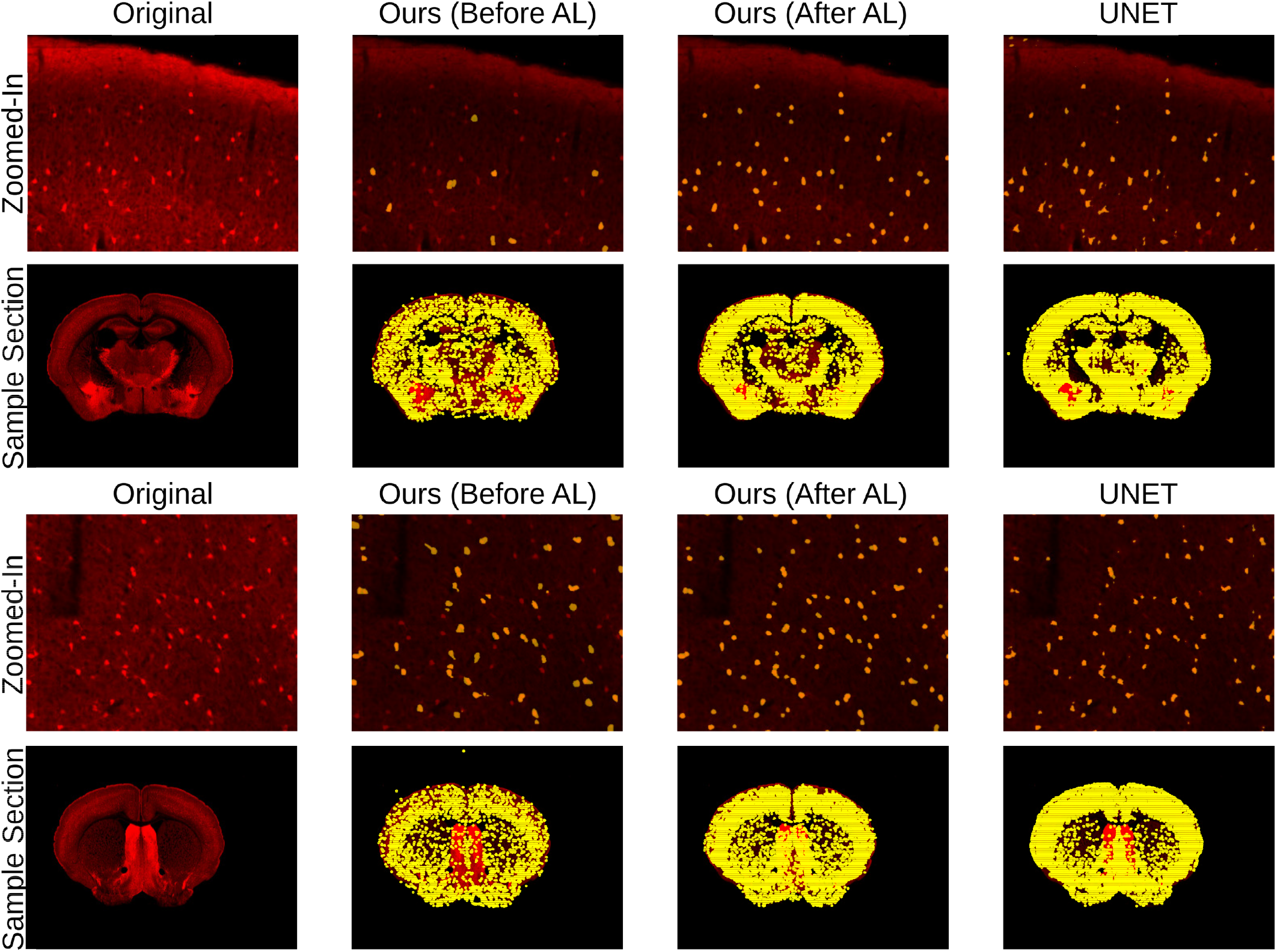
Qualitative results on the unseen Allen Brain dataset. The bottom rows (2nd and 4th) show the original sample, our model’s segmentation mask before Active Learning overlaid, our model’s segmentation masks after Active Learning overlaid, and UNets segmentation masks, from left to right. A zoomed-in subsection, following the same order, is shown in the top rows (1st and 3rd).

Before Active Learning, our model struggled to accurately capture neurons, often producing blob-like masks with a significant number of false positives. However, in post-active learning, our model demonstrates an enhanced capability to precisely identify and generate individual masks for specific neurons. By employing a detection model followed by image segmentation, we effectively address the issue of multiple neurons being amalgamated under a single mask. As illustrated in the zoomed-in subsections in **Figure 2**, our model successfully generates distinct masks for neurons even in close proximity. Additionally, the intensity-based thresholding significantly reduces false positives, resulting in cleaner and more accurate segmentation.

We compare our model’s performance with UNet (Ronneberger et al. [2015b]) using the same data samples. As shown in **Figure 2**, UNet exhibits a higher occurrence of false positives and often misses smaller-sized neurons. The zoomed-in sections reveal that the segmentation masks generated by UNet contain significant broken masks as well.

#### Performance on Fluocells v2 dataset

To establish the baseline performance of our pipeline, we conduct model inference on the Fluocells dataset and compare the results with those obtained using UNet. We utilize the existing test set for qualitative and quantitative evaluation. The pre-processed, normalized images are passed through the neuron detector and segmenter, and the DICE score (Dice [1945]) is computed between the model’s output and the corresponding ground truth. **Figure 3** illustrates the quantitative and qualitative results of our model and UNet on samples corresponding to the three stains present in the dataset. Initially, we observe that our model’s results include the actual neurons (true positives) but also a significant number of false positives, resulting in a low DICE score. To address the issue of excessive false positives, we apply intensity thresholding to the generated masks. As demonstrated in **Figure 3**, this process effectively removed the false positives, leading to a significant increase in the DICE score and producing a cleaner output mask. This improvement was consistently observed across all three stains — cFOS, CTb, and orexin.

**Figure 3:**
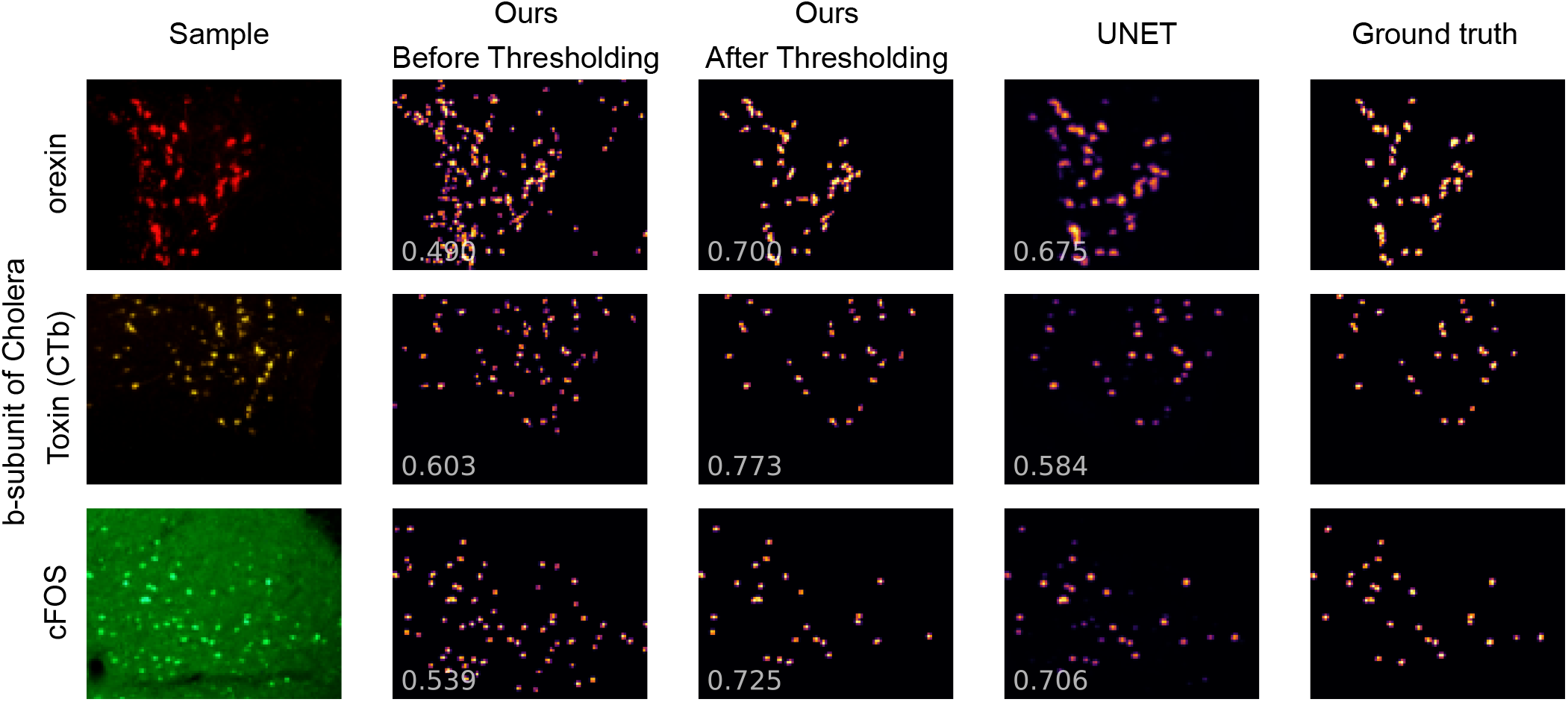
Qualitative results on the unseen Fluocells dataset. The original sample for the 3 stains in the dataset is shown on the left, followed by the masks generated by our framework before intensity thresholding, the masks after applying intensity thresholding, masks generated by UNet, and the actual ground truths. The DICE score between the model’s masks and ground truth is mentioned in the bottom left corner of the masks.

While UNet achieves decent overall performance and a comparable DICE score, our model demonstrates superior performance, particularly in handling samples with varying intensity levels. UNet struggles to capture low-intensity neurons, resulting in missed detections and a lower DICE score in such cases. In contrast, our model is better equipped to handle these variations, leading to more accurate segmentation and higher overall performance.

Using both the Fluocells and SOM-Cre Mouse line datasets, we demonstrate the performance of our neuron segmentation model with and without the application of Active Learning. It is important to note that once the intensity parameter is adjusted, the segmentation results closely match the ground truth, leading to near-perfect ground truth masks.

## Conclusion

We present Segment AnyNeuron, an active learning-based framework for neuron segmentation using Detectron2 and HQ SAM. This approach reduces manual annotation needs by iteratively refining the model with minimal ground truth correction while maintaining high performance. Advanced preprocessing, including intensity normalization and patch-based segmentation, ensures clean inputs and intensity-based thresholding enhances accuracy. We validate our framework on the Fluocells and SOM-Cre mouse line datasets, showing high accuracy and robustness. Active learning on the SOM-Cre dataset further improves performance, avoiding the cold-start problem and optimizing manual annotation of key samples. This framework has significant potential for advancing medical research and drug discovery through precise neuron segmentation.

In conclusion, our framework offers a robust and adaptable solution for neuron segmentation, combining state-of-the-art models with Active Learning efficiency. This method enhances segmentation accuracy and provides a scalable approach for complex medical imaging datasets, paving the way for future innovations in medical image analysis.

## Methods

### Intensity normalization

Medical images, especially fluorescent images, often exhibit varying intensities, posing challenges for DL object detection and image segmentation models. Therefore, our pipeline incorporates essential preprocessing steps, including intensity normalization, to address this issue effectively. The input image is divided into smaller patches that undergo intensity normalization. This process enables efficient handling of high-resolution images, while intensity normalization adjusts the pixel values to significantly reduce overall intensity variability.

The image *I* is split into patches (of size *x*1 x *x*2) and for each patch, an intensity threshold *θ* is calculated which is used for the normalization process.

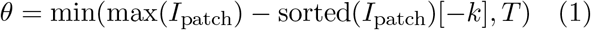

where *k* = 5 and *T* = 10 are constants chosen to identify significant pixel values while avoiding outliers and preventing excessive adjustment. Furthermore, for the purposes of our experiments, we set *x*1 = *x*2 = 256.

Using *θ* and its mean intensity (*µ*), the patch is normalized, followed by gamma correction. The final image patch intensities are then rescaled between the 0.1 and 0.99 percentiles.

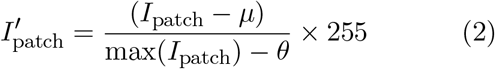

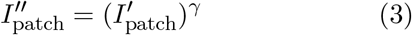

Intensity normalization enhances the contrast of the image leading to a more accurate and robust segmentation. All the intensity-normalized patches are stitched together to reconstruct the original image, which is then fed as input to the neuron detector.

### Neuron Detector

The object detection model we employ as part of our neuron detector is Detectron2, chosen for its widespread use and efficacy in medical image analysis (Salh and Ali [2024], Ali et al. [2022]). Building upon Mask-RCNN (He et al. [2017]), Detectron2 uses a Feature Pyramid Network (FPN) (Lin et al. [2017]) with ResNet (He et al. [2016]) blocks to downsample images and extract hierarchical features. The Region Proposal Network (RPN) (Ren et al. [2015]) processes these features to generate top-scoring bounding boxes, which are refined through the BoxHead for the final output. In our framework, Detectron2 detects neurons in normalized images by using the centers of bounding boxes as keypoints, crucial for accurate neuron identification. While Detectron2 can produce segmentation masks, it is less effective for small neurons, often merging multiple neurons into a single mask. Thus, we rely on the object detection head for precise neuron identification and segmentation. Moreover, during finetuning, we use individual neuron masks within each image to further minimize the possibility of multiple neurons being assigned a single mask. Detectron2 also plays a pivotal role in generating an initial, albeit imperfect, ground truth for our Active Learning pathway. Using keypoints over segmentation masks significantly speeds up the ground truth correction process, as annotating key points is more straightforward and expedient, enhancing annotation efficiency.

### Active Learning

To optimize our model for any neuron data, we employ Active Learning, which allows fine-tuning with minimal ground truth and fewer training iterations. This human-in-the-loop approach involves experts correcting the initial, “dirty” ground truth generated by our neuron detector for the most representative samples. To ensure comprehensive coverage of the feature space during sample selection for manual annotation, we embed the dataset into a low-dimensional space using Uniform Manifold Approximation and Projection (UMAP) (McInnes et al. [2020]. We then select representative samples from both sparse and dense clusters for use in the active learning pipeline.

Starting with the neuron detector’s output provides an initial baseline, reducing manual labeling effort and circumventing the cold-start problem commonly associated with Active Learning (Chen et al. [2022]). This iterative process of refining the ground truth and continuously updating the model enhances its generalization capabilities and enables rapid convergence to a highly accurate state.

### Neuron Segmenter

The refined keypoints from the Active Learning step are fed into the Neuron segmenter. We use HQ SAM, a state-of-the-art segmentation model as part of our pipeline as it excels at processing keypoints to generate high-quality segmentation masks, even in complex and noisy images, achieving precise neuron segmentation. The process begins with the refined keypoints, which are corrected through minimal ground truth annotation during the Active Learning phase. These keypoints serve as crucial landmarks, guiding HQ SAM to focus on specific regions of interest within the image patches. The accurate reference points provided by these keypoints significantly enhance the precision of the segmentation. This approach helps mitigate issues of overlapping and closely packed neurons.

The precise masks generated by HQ SAM are more reliable and accurate. Once the masks are generated, intensity-based thresholding is applied to filter out low-intensity neurons, enhancing the overall segmentation accuracy.

### Intensity-based thresholding

We apply intensity-based thresholding to the masks generated by HQ-SAM since it allows us to filter out low-intensity neurons, which are often false positives. By adjusting the intensity threshold, users can control the inclusion of neurons in the final segmentation mask, optimizing the results based on their specific requirements. The user-controlled intensity knob provides flexibility and customization, ensuring that the segmentation meets the desired accuracy and specificity. Post-segmentation, the intensity of each neuron is measured using the original image to ensure accurate intensity values. These values are then normalized for consistency. Users can adjust the intensity threshold, which allows them to filter out neurons that do not meet the desired intensity criteria. This step facilitates in removing false positives and improving the overall accuracy of the segmentation by giving users the freedom to exclude neurons based on their specific intensity requirements. This flexibility is essential for tailoring the segmentation to different applications and datasets, enhancing the framework’s effectiveness. To quantify our model’s performance, we compare our results with U-Net (Ronneberger et al. [2015c]), a well-established model in medical image segmentation. U-Net serves as a benchmark in the domain, particularly for medical datasets, and is widely used by recent models to demonstrate segmentation efficacy (Weng et al. [2019], Cao et al. [2022]). Its architecture, featuring a contracting path for context capture and an expansive path for precise localization, makes it exceptionally effective for tasks such as neuron segmentation. U-Net’s ability to work well with limited annotated data and produce high-resolution segmentation maps has led to its widespread adoption and significant success in various medical imaging applications. This makes it an ideal model for benchmarking and comparing new segmentation algorithms (Azad et al. [2022]). Prior to experimentation, we trained UNet on the pre-defined training set of Fluocells for approximately 100 epochs and used that checkpoint for performance comparisons. We demonstrate that our model performs marginally better than UNet on both the Fluocells and SOM-Cre Mouse Line datasets.

## References

Yuxin Wu, Alexander Kirillov, Francisco Massa, Wan-Yen Lo, and Ross Girshick. Detectron2. https://github.com/facebookresearch/detectron2, 2019.

Lei Ke, Mingqiao Ye, Martin Danelljan, Yifan Liu, Yu-Wing Tai, Chi-Keung Tang, and Fisher Yu. Segment anything in high quality. In NeurIPS, 2023.

Shervin Minaee, Yuri Boykov, Fatih Porikli, Antonio Plaza, Nasser Kehtarnavaz, and Demetri Terzopoulos. Image segmentation using deep learning: A survey. IEEE transactions on pattern analysis and machine intelligence, 44(7):3523–3542, 2021.

Olaf Ronneberger, Philipp Fischer, and Thomas Brox. U-net: Convolutional networks for biomedical image segmentation, 2015a. URL https://arxiv.org/abs/1505.04597.

Sudhanshu Mittal, Joshua Niemeijer, Jörg P. Schäfer, and Thomas Brox. Best practices in active learning for semantic segmentation, 2023. URL https://arxiv.org/abs/2302.04075.

Radek Mackowiak, Philip Lenz, Omair Ghori, Ferran Diego, Oliver Lange, and Carsten Rother. Cereals - cost-effective region-based active learning for semantic segmentation, 2018. URL https://arxiv.org/abs/1810.09726.

Chieh-Chi Kao, Teng-Yok Lee, Pradeep Sen, and Ming-Yu Liu. Localization-aware active learning for object detection, 2018. URL https://arxiv.org/abs/1801.05124.

Vishwesh Nath, Dong Yang, Bennett A. Landman, Daguang Xu, and Holger R. Roth. Diminishing uncertainty within the training pool: Active learning for medical image segmentation. IEEE Transactions on Medical Imaging, 40(10): 2534–2547, October 2021. ISSN 1558-254X. doi: 10.1109/tmi.2020.3048055. URL 10.1109/TMI.2020.3048055.

Andrew S Boehringer, Amirhossein Sanaat, Hossein Arabi, and Habib Zaidi. An active learning approach to train a deep learning algorithm for tumor segmentation from brain mr images. Insights into Imaging, 14(1):141, 2023.

Asim Iqbal, Asfandyar Sheikh, and Theofanis Karayannis. Denerd: high-throughput detection of neurons for brain-wide analysis with deep learning. Scientific reports, 9(1):13828, 2019.

Chiman Haydar Salh and Abbas M Ali. Automatic detection of breast cancer for mastectomy based on mri images using mask r-cnn and detectron2 models. Neural Computing and Applications, 36 (6):3017–3035, 2024.

Ammar Alhaj Ali, Rasin Katta, Roman Jasek, Bronislav Chramco, and Said Krayem. Covid-19 detection from chest x-ray images using detectron2 and faster r-cnn. In Proceedings of the Computational Methods in Systems and Software, pages 37–53. Springer, 2022.

Farheen Chincholi and Harald Koestler. Detectron2 for lesion detection in diabetic retinopathy. Algorithms, 16(3):147, 2023.

Yichi Zhang, Zhenrong Shen, and Rushi Jiao. Segment anything model for medical image segmentation: Current applications and future directions. Computers in Biology and Medicine, page 108238, 2024.

Cécile Viollet, Axelle Simon, Virginie Tolle, Alexandra Labarthe, Dominique Grouselle, Yann Loe-Mie, Michel Simonneau, Guillaume Martel, and Jacques Epelbaum. Somatostatin-ires-cre mice: between knockout and wild-type? Frontiers in endocrinology, 8:131, 2017.

Declan Williams, Bei Qi Yan, Hansen Wang, Logine Negm, Christopher Sackmann, Claire Verkuyl, Vanessa Rezai-Stevens, Shehab Eid, Nimit Vediya, Christine Sato, et al. Somatostatin slows aβ plaque deposition in aged app nl-f/nl-f mice by blocking aβ aggregation. Scientific Reports, 13(1):2337, 2023.

Fadi Rofo, Friederike A Sandbaumhuter, Aikaterini Chourlia, Nicole G Metzendorf, Jamie I Morrison, Stina Syvanen, Per E Andrén, Erik T Jansson, and Greta Hultqvist. Wide-ranging effects on the brain proteome in a transgenic mouse model of alzheimer’s disease following treatment with a brain-targeting somatostatin peptide. ACS Chemical Neuroscience, 12(13):2529–2541, 2021.

Olaf Ronneberger, Philipp Fischer, and Thomas Brox. U-net: Convolutional networks for biomedical image segmentation. In International Conference on Medical Image Computing and Computer-Assisted Intervention, pages 234–241. Springer, 2015b.

Lee R Dice. Measures of the amount of ecologic association between species. Ecology, 26(3):297–302, 1945.

Kaiming He, Georgia Gkioxari, Piotr Dollár, and Ross Girshick. Mask r-cnn. In Proceedings of the IEEE International Conference on Computer Vision (ICCV), 2017.

Tsung-Yi Lin, Piotr Dollár, Ross Girshick, Kaiming He, Bharath Hariharan, and Serge Belongie. Feature pyramid networks for object detection. In Proceedings of the IEEE Conference on Computer Vision and Pattern Recognition (CVPR), 2017.

Kaiming He, Xiangyu Zhang, Shaoqing Ren, and Jian Sun. Deep residual learning for image recognition. In Proceedings of the IEEE Conference on Computer Vision and Pattern Recognition (CVPR), pages 770–778, 2016.

Shaoqing Ren, Kaiming He, Ross Girshick, and Jian Sun. Faster r-cnn: Towards real-time object detection with region proposal networks. In Proceedings of the 28th International Conference on Neural Information Processing Systems (NIPS), pages 91–99, 2015.

Leland McInnes, John Healy, and James Melville. Umap: Uniform manifold approximation and projection for dimension reduction, 2020. URL https://arxiv.org/abs/1802.03426.

Liangyu Chen, Yutong Bai, Siyu Huang, Yongyi Lu, Bihan Wen, Alan L. Yuille, and Zongwei Zhou. Making your first choice: To address cold start problem in vision active learning, 2022. URL https://arxiv.org/abs/2210.02442.

Olaf Ronneberger, Philipp Fischer, and Thomas Brox. U-net: Convolutional networks for biomedical image segmentation, 2015c. URL https://arxiv.org/abs/1505.04597.

Yu Weng, Tianbao Zhou, Yujie Li, and Xiaoyu Qiu. Nas-unet: Neural architecture search for medical image segmentation. IEEE access, 7:44247–44257, 2019.

Hu Cao, Yueyue Wang, Joy Chen, Dongsheng Jiang, Xiaopeng Zhang, Qi Tian, and Manning Wang. Swin-unet: Unet-like pure transformer for medical image segmentation. In European conference on computer vision, pages 205–218. Springer, 2022.

Reza Azad, Ehsan Khodapanah Aghdam, Amelie Rauland, Yiwei Jia, Atlas Haddadi Avval, Afshin Bozorgpour, Sanaz Karimijafarbigloo, Joseph Paul Cohen, Ehsan Adeli, and Dorit Merhof. Medical image segmentation review: The success of u-net, 2022. URL https://arxiv.org/abs/2211.14830.

